# Analysis of Virus-Host Interactomes through a network-centric approach

**DOI:** 10.1101/2020.06.30.179945

**Authors:** Giuseppe Tradigo, Pierangelo Veltri, Pietro Hiram Guzzi

**Affiliations:** DEPARTMENT OF MEDICAL AND SURGICAL SCIENCES, UNIVERSITY OF CATANZARO

**Keywords:** protein-to-protein interaction, interactome, virus, graph theory

## Abstract

Viruses are small microorganisms that invade living cells and use them to replicate themselves. Viruses cause many common human infections (such as cold flu) as well as many lethal diseases. Therefore the comprehensive knowledge of mechanisms used by viruses to infect living cells, also known as host-pathogen interactions, is crucial. Mechanisms of infections of viruses are mediated by Protein-protein interactions (PPIs). PPIs are often modelled using graphs, thus the use of such a model may also explain mechanisms of infection of viruses. In this work, we propose a methodology to model and analyse host-pathogen interactions and a supporting tool able to analyse such data. We also analyse host-pathogen interactions of some common viruses demonstrating common mechanism and differences.

## 1. Introduction

Viruses are small microorganisms which use living cells to replicate themselves. Viruses cause many infectious diseases that are responsible for millions of death every year [18]. They exist in the form of small independent particles named virions. Each virion consists of two main components: (*i*) the genetic information, encoded as DNA or RNA and (*ii*) a protein coat, named capsid, which wraps the genetic material. Sometimes the capsid is surrounded by an envelope of lipids. Virions have different shapes that are used for classification of themselves [7].

Viruses are not able to replicate themselves alone, therefore they must use the metabolism of an **host** organism to reproduce themselves. The virus replication cycle may be summarised into six main steps [15]:

**(1) Attachment**. First, viruses bind the surface of host cells.

**(2) Penetration**. Viruses enter the host cell through receptor-mediated endocytosis or membrane fusion.

**(3) Uncoating**. The viral capsid is removed and virus genomic materials are released.

**(4) Replication**. Viruses use the host cells to replicate their genomic information. In this step viral proteins are synthesized and possibly assembled. Viral proteins may interact among them and with host proteins to perform their function (e.g. regulate the protein expression).

**(5) Assembly**. Following the structure-mediated self-assembly of the virus particles, some modifications of the proteins often occur.

**(6) Release**. Viruses can be released from the host cell by lysis, a process that kills the cell.

During the replication step, proteins of the virus use the host environment, interact among them and with the host proteins, causing loss of function or even the death of the cells. Therefore the complete elucidation of the whole set of such interactions is a crucial step for the comprehension of viruses.

Nowadays, thanks to the use of different proteomic technologies, the complete set of interactions is available for many viruses [10, 4, 12].

Proteins of viruses may interact among them (virus-virus host interactions), or may interact with proteins of the host organism (virus-host interactions). Such interactions are usually modeled by using graphs and stored in a growing number of databases such as: Virus Mint [8], String Viruses [9], HpiDB [2], Virus Mentha [3], and VirHostNet [14].

The importance of understanding the interplay between host and virus proteins is relevant since it may help in identifying vires-related diseases as well as potential targets for therapeutic strategies. Consequently, some studies presented the investigation of virus-host interactomes using tools and methodologies coming from graph theory [19, 4], demonstrating the importance of studying virus-host interplay at network level [21, 5, 16, 11].

For instance [20] Uetz et al., analysed the interactome of some viruses on a network scale. They computed some network characteristics such as **diameter, characteristic path length, local clustering coefficient**, and similarity with respect to the scale free model. They found that network characteristics of such viruses are different from other species. Then they merged the virus interactome with the host one concluding that merged interactome present similar characteristics of the host interactome alone.

Vidal et al., [11] presented probably the first work describing the characteristics of human proteins attached by virus in terms of network properties. Authors integrate data from all the database at the date of publication (year 2007), and they conclude that human proteins that interact with viral ones are preferably hub or bottleneck [6, 1].

Research in this field is based on the availability of reliability of data. Nowadays there exist lots of data with respect to those available in previous studies, e.g. the HpiDB [2] contains more than 40.000 interactions while many previous studies are usually based on the analysis of less than one thousands interaction. Consequently there is still room for novel research projects providing both novel methodologies as well as supporting tools.

Main characteristics of such works are: the study of a single virus, and the analysis of network properties. From a bioinformatic point of view, there is still room for the investigation and comparison of different viruses and the modification in the host interactomes in a single framework, as well as the lack for a unified software framework enabling such studies.

Consequently, we here propose a bioinformatic methodology aiming at the investigation of such relevant questions: are the proteins infected by viruses central or peripheral (i.e. are the infected proteins hub or not)?; do all of the viruses attach to similar proteins (from a network point of view)?; what happens in an infected host interactome?

Literature reports that interactomes usually have common properties [13]: a modular organization, a small-world property (i.e. great connectivity between proteins), the presence of *communities*, and some more *relevant* proteins, i.e. more central proteins, also referred to as hubs.

Some have argued that these central proteins, or hubs, are essential to biological functions. In this study, we want to explore and compare the centrality of host proteins attached by viruses.

We also propose a software tool able to import and analyse virus and host data enabling the user to easily investigate such properties: (i) network centrality of infected proteins, (ii) modification of host interactomes, (iii) comparison of different interactomes.

## 2. Network Centrality Measures

A network is represented by a graph *G* = (*V, E*), where *V* is a set of nodes and *E* ∈ *V* ×*V* is a set of (ordered) pairs of nodes called edges. A graph is usually represented by using **g** ∈ ℜ^*n*×*n*^, where *g*_*ij*_ ≠ 0 indicates the existence of an edge between nodes *i* and *j* and *g*_*ij*_ ≠ 0 indicates the absence of an edge between the two nodes. A special case of graphs, called **edge-weighted graphs**, is characterized by a special adjacency matrix whose values are real valued values included in the interval (0, 1). The following discussion is focused on undirected non-weighted graphs and may be easily extended to both ordered and edge-weighted graphs.

The *degree* of a node *i* in a undirected graph *G*, denoted *deg*_*i*_(**g**) = |{*j*: *g*_*ij*_ ≠ 0}|, is the number of nodes adjacent to *i* Two nodes *w*_*i*_ and *w*_*j*_ are connected (or path-connected) if there exists a path between them.

In the case of an unweighted network, a *geodesic* (or shortest path) from node *w*_*i*_ to node *w*_*j*_ is the path the involves the minimum number of edges. Consequently, we may define the *distance* between nodes *w*_*i*_ and *w*_*j*_, where *ρ*_**g**_(*w*_*i*_, *w*_*j*_) is the number of edges involved in a geodesic between *w*_*i*_ and *w*_*j*_.

Starting from the computation of distance, a set of *centrality* measures have been introduced. The aim of such measure is to evidence the relevance, or importance, of a node in a network by analyzing the topology.

The simplest measure is the **Degree centrality**. Given a node *w*_*i*_, the degree centrality is the number of adjacent nodes to *w*_*i*_, defined as

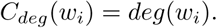

Sometimes, the degree centrality is normalized by the maximal possible degree of a node, to obtain a number between 0 and 1:

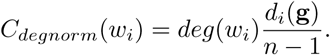

Degree centrality is an obvious measures and its computation is also simple. It gives some information related to the relevance of the node *w*_*i*_ but it misses some relevant aspects of the whole structure of the network as well as the node’s position [**?**].

Other centrality measures are based on calculation of the geodesic paths and on the analysis of the distance of a node with respect to the others. The rationale of the **Closeness centrality** is to consider as central a node that is *close* to the others in terms of distance.

Formally, the closeness centrality of a node *w*_*i*_ is the reciprocal of the average shortest path distance to *w*_*i*_ over all n-1 reachable nodes, i.e.

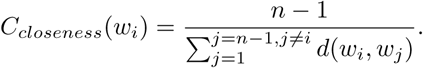

where *d*(*w*_*i*_, *w*_*j*_) is the shortest distance among *w*_*i*_ and *w*_*j*_. Closeness centrality has a simple calculation and it measures how a node is reachable by the others. In such formulation each connected component of a graph present a different closeness centrality distribution. In order to overcome this limitation, Wasserman and Faust propose an improved formula for graphs with more than one connected component [22].

The closeness centrality indicates how a node is close to the other, while sometimes it is important to evaluate how a node stands between each other. For these aims the **Betweenness centrality** has been introduced [17]. Betweenness centrality first calculates all the shortest paths among all node pairs. Then, for each node *w*_*i*_, the number of shortest paths that pass through *w*_*i*_ is calculated and each node receives a score based on the number of these shortest paths that pass through. Consequently, nodes that are included most frequently into these shortest paths will get a higher betweenness centrality scores.

Formally, the betweenness centrality of a node (*w*_*i*_)

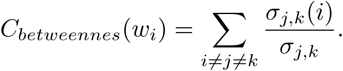

where *σ*_*j,k*_ is the total number of shortest paths from node *w*_*j*_ to node *w*_*k*_ and *σ*_*j,k*_(*i*) is the number of those paths that pass through *i*.

**Eigenvector centrality**, proposed by [**?**], is a measure of the influence of a node in a network. It scores all nodes of a network on the assumption that connections to high-scoring nodes contribute more to the score of the node rather than connections to low-scoring nodes.

## 3. Host-Pathogen data

The VirusMINT database [8] is derived from the MINT database and it is based on the collection of protein interaction data from literature. It currently stores over 5000 interactions including both virus-virus and virus host interactions. It offers a simple graphical query interface available through a web browser (http://mint.bio.uniroma2.it/virusmint/).

Virus Mentha [3] collects information about viral interaction from many sources (e.g. experiments, literature). It also offers some analysis capabilities on a network level as well as a reliability score for each interaction which takes into account all of the supporting evidence. String Viruses [9] combines evidence from experimental and text-mining channels to provide combined probabilities for interactions between viral and host proteins. The database contains 177,425 interactions between 239 viruses and 319 hosts. VirHostNet [14] contains both virus–virus and virus–host protein–protein interactions as well as their projection onto their corresponding host cell protein interaction networks.

The HpiDB database [2] contains 45.238 manually curated entries in the current release. It stores 594 pathogen and 70 host species. It also offers to the users additional Gene Ontology functional information and an implementation of network visualization.

## 4. Analysis of Host-Pathogen Interactomes

Vidal et al., [11] presented probably the first work describing the characteristics of human proteins attached by virus in terms of network properties. Authors integrate data from all the database at the date of publication (year 2007), and they conclude that human proteins that interact with viral ones are preferably hub or bottleneck. Neverthless, data related to host interactomes are regularly growing due to both wet and in silico experiments, therefore there is the need to perform novel analyses.

Therefore we propose a novel general methodology for such analyses based on following steps, as depicted in Figure 1:

**Figure 1.**
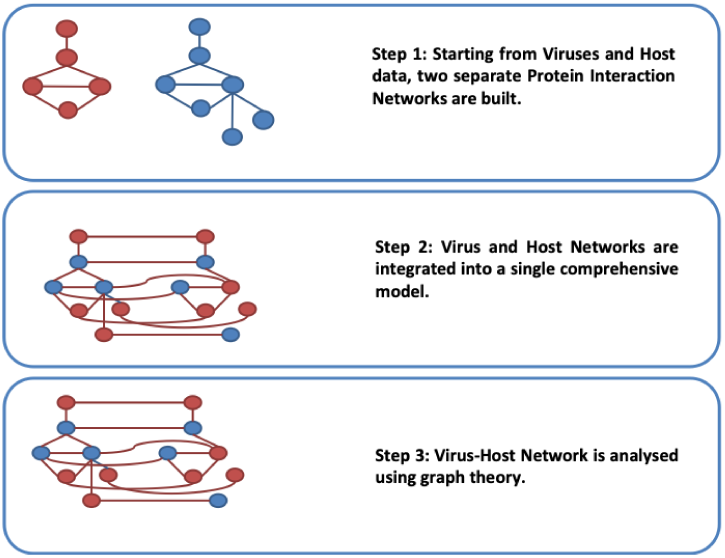
Our Methodology.

1. generate two separate protein interaction networks for virus and host organisms starting from available data;
2. integrate virus and host networks into a single comprehensive model;
3. calculate centrality for all the proteins of the host organism and we compare centrality of attached proteins with respect to the non attached proteins.

## 5. VirnetAnalyzer: A TOOL FOR THE AUTOMATED ANALYSIS OF THE HOST-PATHOGEN INTERACTOMES

We designed VirNetAnalyzer, a tool for the automated analysis of the centrality of proteins of an host organism attached by a virus. We also implemented a preliminary prototype, as depicted in Figure 2, using the Python programming language. The software tool is able to read data of virus and host interactomes. Once data has been loaded, it calculates the centrality of host proteins attached by viruses and it compares these values with the median values calculated for the host. It also includes a visualization module able to represent the host interactome. The software is free for academic purposes and it is available upon request.

**Figure 2.**
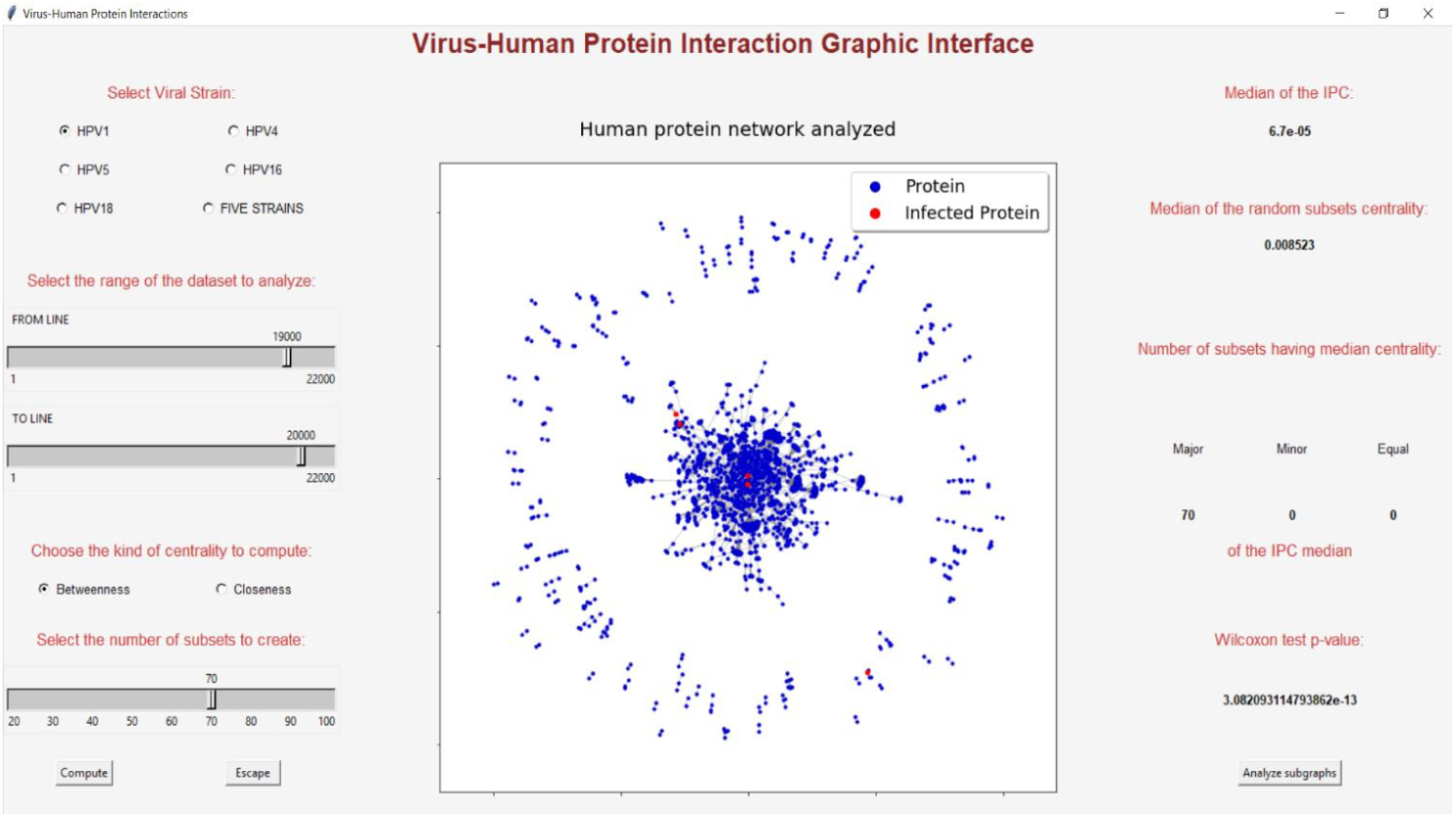
The GUI of VirNetAnalyzer.

## 6. Conclusion

In this work we proposed a methodology to model and analyse host-pathogen interactions and a supporting tool able to analyse such data. We analyzed host-pathogen interactions of some common viruses demonstrating both common mechanisms and differences.

## Footnotes

1 Authors are listed in alphabetical order

